# The Impact of the Local Mechanical Environment on Cell Shape and Chondrogenesis of Mesenchymal Stromal Cells in 3D Biomimetic Composite Materials

**DOI:** 10.1101/2023.08.30.555478

**Authors:** M. Fenu, I. Muntz, D. Harting, J. Xu, M. D’Este, G.H. Koenderink, G.J.V.M van Osch

## Abstract

Efforts to model and repair connective tissue through engineered tissue constructs have generated great interest in culturing cells in 3d polymer network environments. It has been shown that the polymer environment is influential in determining cellular responses such as differentiation, migration and morphology. Hydrogels are used to mimic the cellular microenvironment, but in most cases hydrogels consisting of one polymeric component are used whereas tissues are composites of different polymers. A clear understanding of how different extracellular components and their mechanical characteristics influence cell behaviour is lacking. Here we developed and characterised composite hydrogels of hyaluronan and fibrin and evaluated their use for cartilage tissue engineering. We demonstrate that these cartilage-mimicking composites have a higher stiffness relative to the individual constituents. Next, we cultured human mesenchymal stromal cells in these 3D hydrogels with chondrogenic media and revealed marked differences in cell morphology, gene expression and cartilage-like matrix deposition depending on the specific extracellular composition. We found that, despite evidence for strong adhesion of the cells to fibrin networks in 2D systems, in 3D systems the primary determinant of cellular morphology is the significantly denser hyaluronan network. Dense hyaluronan hydrogels cause local cell confinement evidenced by rounder cell morphologies, independent of the presence of fibrin. While the composite fibrin-hyaluronan hydrogels led to lower expression of chondrogenic genes than hyaluronan alone, the larger linear modulus and resistance to cell-mediated contraction due to the composite nature of the matrix provides a strong advantage in terms of macroscopic mechanical stability. These findings highlight the potential of multi-component hydrogels for controlling cellular behaviour and bulk mechanical properties of cell-hydrogel constructs independently, therefore opening avenues for better understanding the complex interplay between cells and their extracellular environment and thus improve the biofabrication of connective tissues for disease modelling and tissue regeneration.

**Graphical Abstract:** 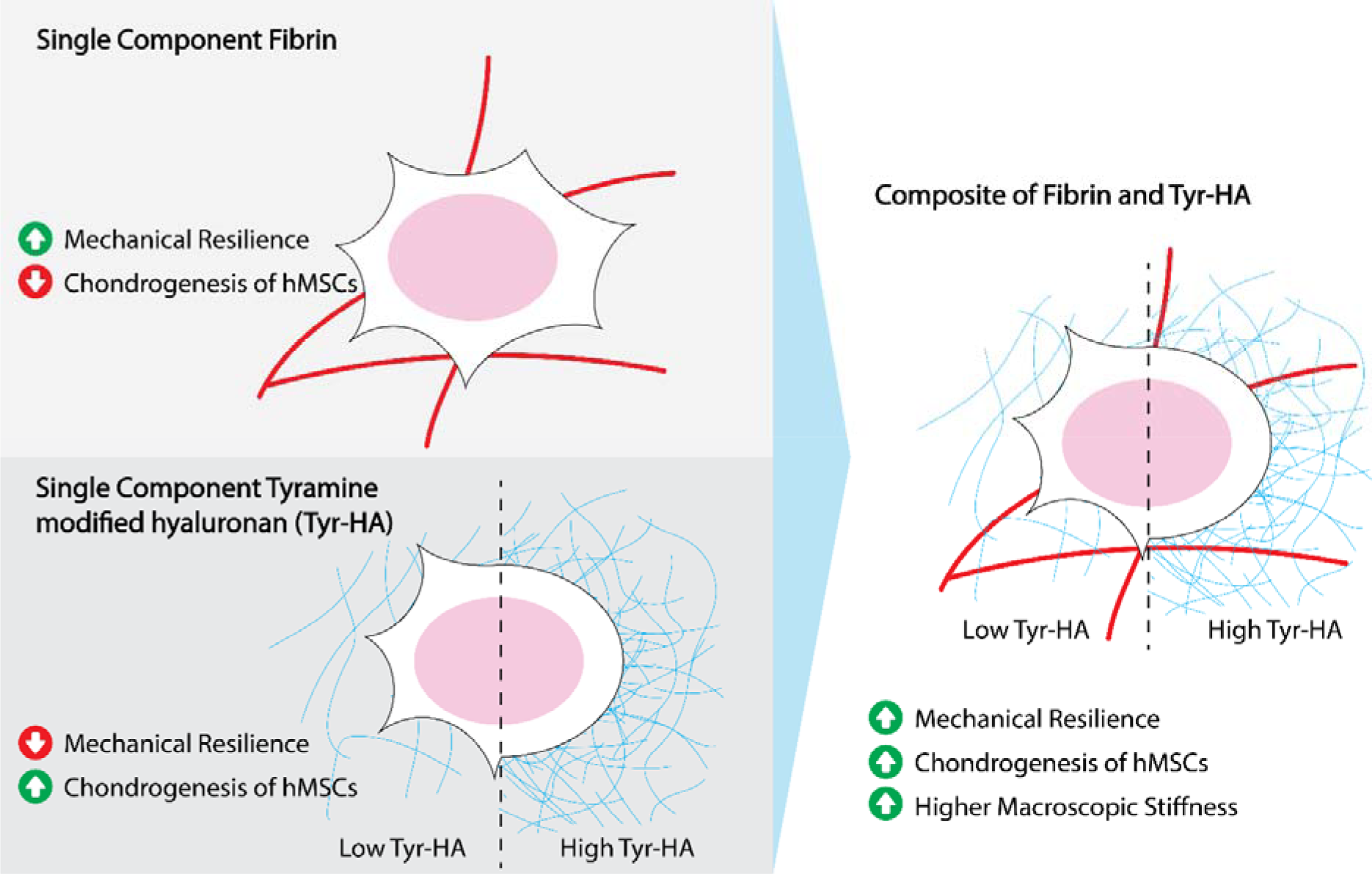

## 1. Introduction

Interest in hydrogels for cell culture has grown rapidly in recent years, driven by the recognition that hydrogels can provide mechanical and biological conditions that mimic tissue-specific properties [1]. Biomimetic hydrogels therefore provide a useful class of cell culture materials for a wide variety of biofabrication and tissue engineering applications, ranging from generation of *in vitro* human disease models to *in vivo* regenerative medicine [2–4]. Hydrogels are a class of materials that have a high water content, similar to natural tissues [5]. Common choices for tissue engineering are materials whose primary components are natively found in the human body, such as fibrin [6, 7], collagen [8] or hyaluronic acid (HA) [9, 10]. In addition to their inherent biocompatibility, these materials provide mechanical and structural integrity. Fibrin has especially advantageous mechanical properties because it forms hydrogels that strongly strain-stiffen, helping to protect materials from rupture [11–13]. While HA lacks the stiffening property which fibrin has, it has the advantage that it can be tuned to form hydrogels with different properties (mechanics, permeability and structure) by varying the molecular weight, concentration, or chemical functionalisation [10, 14, 15]. This tunability is important because both the physical and biological functionality of hydrogels strongly influence the cellular response. The viscoelastic properties of hydrogels influence cell spreading and cell differentiation [1], the hydrogel structure influences cell behaviour through porosity and dimensionality [16, 17], and the biological functionality influences cells by signalling to different adhesion receptors [18].

Despite these advances in designing biomimetic hydrogels for biofabrication and tissue engineering, we are still far from modelling the full physical and biological complexity of the native extracellular environment of cells. The hydrogels used mostly consist of one polymeric component. The native extracellular matrix (ECM), however, is composed of many proteins, polysaccharides, and proteoglycans. This complex composition renders the extracellular matrix multifunctional and tailored for each tissue and organ [19]. The matrix at the same time provides high macroscopic mechanical stability to tissues and local tissue-specific physical and biochemical cues to the resident cells. It is challenging to achieve such a complex combination of properties in single-component hydrogels, and it is also difficult to independently tune all these properties using just a single polymer. One solution to tackle this challenge is to use composite hydrogels of two or more biopolymers that offer complementary properties that can be independently tuned. For example, an interpenetrating hyaluronan and fibrin network has been shown to allow the construct to maintain its shape in systems where a pure fibrin network becomes completely degraded [20]. Hyaluronan has also been combined with collagen to improve the printability of the tissue constructs in 3D printing technologies, which allows control over features from macroscopic tissue shape down to microscopic alignment of tissue components [21].

A challenge of tissue engineering remains to design systems with biophysical properties which closely resemble those of native tissues, while providing the appropriate mechanical and biochemical cues to allow desired cellular behaviours such as differentiation and proliferation. One hurdle is the lack of understanding of the physical properties of composite hydrogels, which are composed of two or more materials. Despite a limited understanding of the underlying mechanisms, HA composite materials have been shown to be supportive of chondrogenesis and cartilage-like matrix deposition, which is true for different HA modifications including tyramine modified HA [22, 23]. Cartilage is well suited as a model tissue due to its high matrix-to-cell ratio (>10) [24], its unique mechanical and biological properties and its potential applications in tissue engineering due to the tissue’s limited self-healing capacity [25]. Previous work evaluating the combination of hyaluronan with fibrous proteins, such as collagen, have linked the incorporation of a fibrous network in hyaluronan-based hydrogels to enhanced mechanical properties (i.e., higher shear moduli)[26]. Additionally, these materials have demonstrated good compatibility with both chondrocytes and mesenchymal stromal cells (MSCs) [23, 27]. Yet, the underlying mechanisms of action and, therefore, the principles for designing optimal biomaterials as tissue substitutes remain elusive.

Here we explore the impact of composite biopolymer hydrogels on cell morphology and chondrogenesis of hMSCs using a composite system of fibrin and cross-linked tyramine modified HA (Tyr-HA). We show that combining fibrin and hyaluronan results in composite materials with advantageous synergistic macroscopic mechanical properties and that the material at the same time promotes chondrogenic differentiation of human mesenchymal stromal cells (hMSCs). We find that, while the fibrin network is essential to prevent macroscopic cell-mediated contraction of the 3D hydrogel, the cellular behaviour is primarily governed by the finer Tyr-HA network. We propose that the confinement of the cells by the finer hyaluronan meshwork is the primary contributor to cellular morphology rather than the formation of cell-matrix adhesion points.

## 2. Results

### 2.1 Fibrin Prevents Cell-Induced Contraction in Composite Hydrogels

To prepare hydrogels that combine the unique properties of fibrous protein and glycosaminoglycan (GAG) present in the extracellular matrix of connective tissues, we prepared composite hydrogels of fibrin and Tyr-HA. We focussed on the impact of the hydrogel composition on hMSCs, given the known capability of hMSCs to respond to mechanical stimuli [28] and their widely studied potential in tissue engineering of cartilage [29]. We cultured the constructs and analysed their macroscopic appearance after 14 days (Figure 1). Pure Tyr-HA hydrogels with concentrations of 2 mg/mL, 4 mg/mL, and 8 mg/mL showed significant cell-induced shrinkage. Shrinkage of the constructs gradually decreased with increasing Tyr-HA concentration; the constructs maintained 21.4% of the original diameter at 2 mg/mL to 88.3% at 8 mg/mL. In contrast, composites that combined 2 mg/ml fibrin and Tyr-HA exhibited no discernible cell-mediated shrinkage (see supplementary table 1 for detailed statistical analysis). This effect cannot be attributed to the higher biopolymer concentration alone, and it is likely due to the fibrin network, since the pure fibrin networks also did not show cell-mediated shrinkage. These findings suggest that the presence of fibrin in the hydrogel matrix contributes to the macroscopic mechanical integrity under the influence of cell-generated forces.

**Figure 1:**
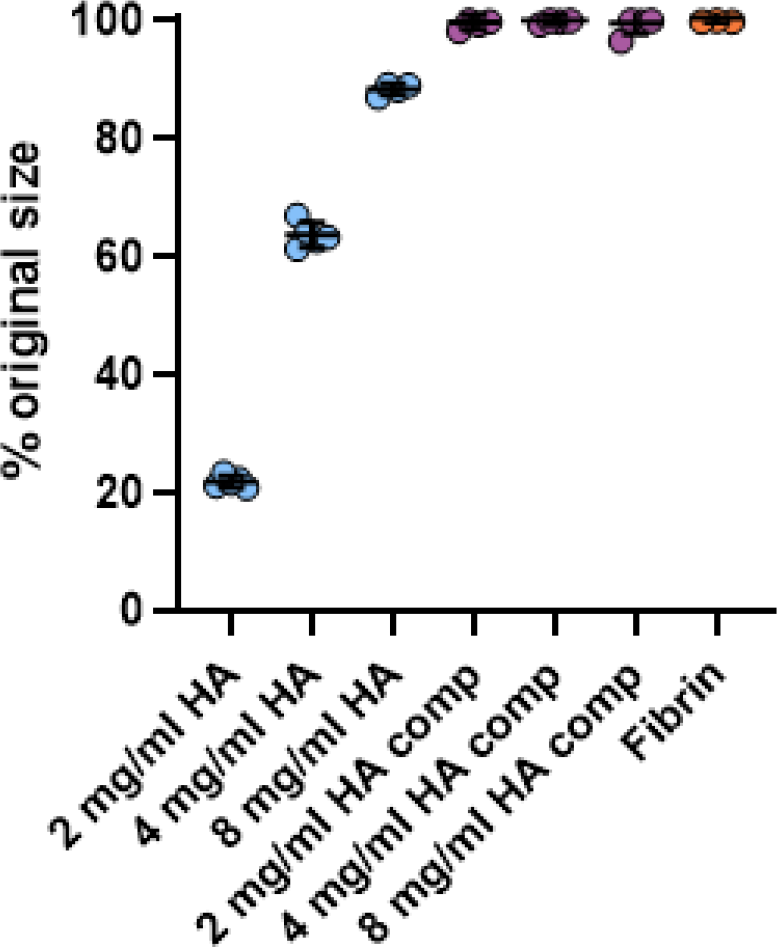
The presence of fibrin in hydrogels with tyramine modified hyaluronan (Tyr-HA) prevents macroscopic cell-induced gel contraction. Average diameter of the constructs after 14 days of culture of hMScs at a seeding density of 3*10 cells/ml normalised by the original diameter. Hydrogels were composed of 2 mg/ml, 4 mg/ml or 8 mg/ml Tyr-HA, with or without an additional 2 mg/ml fibrin (composites with fibrin are denoted as “comp”). Data for cells in pure fibrin are shown on the right. Data are presented as the mean and standard deviation of 5 samples, with each point representing a separate sample.

### 2.2 Modulating Hyaluronan Concentration Enhances Mechanical Properties of Composite Hydrogels

All hydrogels showed strain-stiffening behaviour (figure 2(a)). The fibrin networks stiffened at much smaller strain (∼10 %) than the Tyr-HA networks (∼100 %), reflecting the much larger persistence length of fibrin fibres compared to hyaluronan polymers [30–32]. Interestingly, the fibrin/Tyr-HA composites exhibited a strong enhancement of the linear modulus when compared to the individual constituents (Figure 2(a)). For composites with low Tyr-HA concentrations (2 mg/mL or 4 mg/mL), the absolute magnitude of the modulus was 10 times larger than for the pure fibrin network and pure 4 mg/ml Tyr-HA network and 1000 times larger than for the 2 mg/ml Tyr-HA network. At the same time, the moduli at these concentrations were also larger than the sum of the moduli of the constituents. For the 8 mg/ml Tyr-HA composite, the stiffness at low strains was comparable to that of the pure Tyr-HA system, but there remained an enhancement of the shear modulus at high strains, where the stiffness reached values at least 10 times larger than for either the pure Tyr-HA or the pure fibrin.

**Figure 2:**
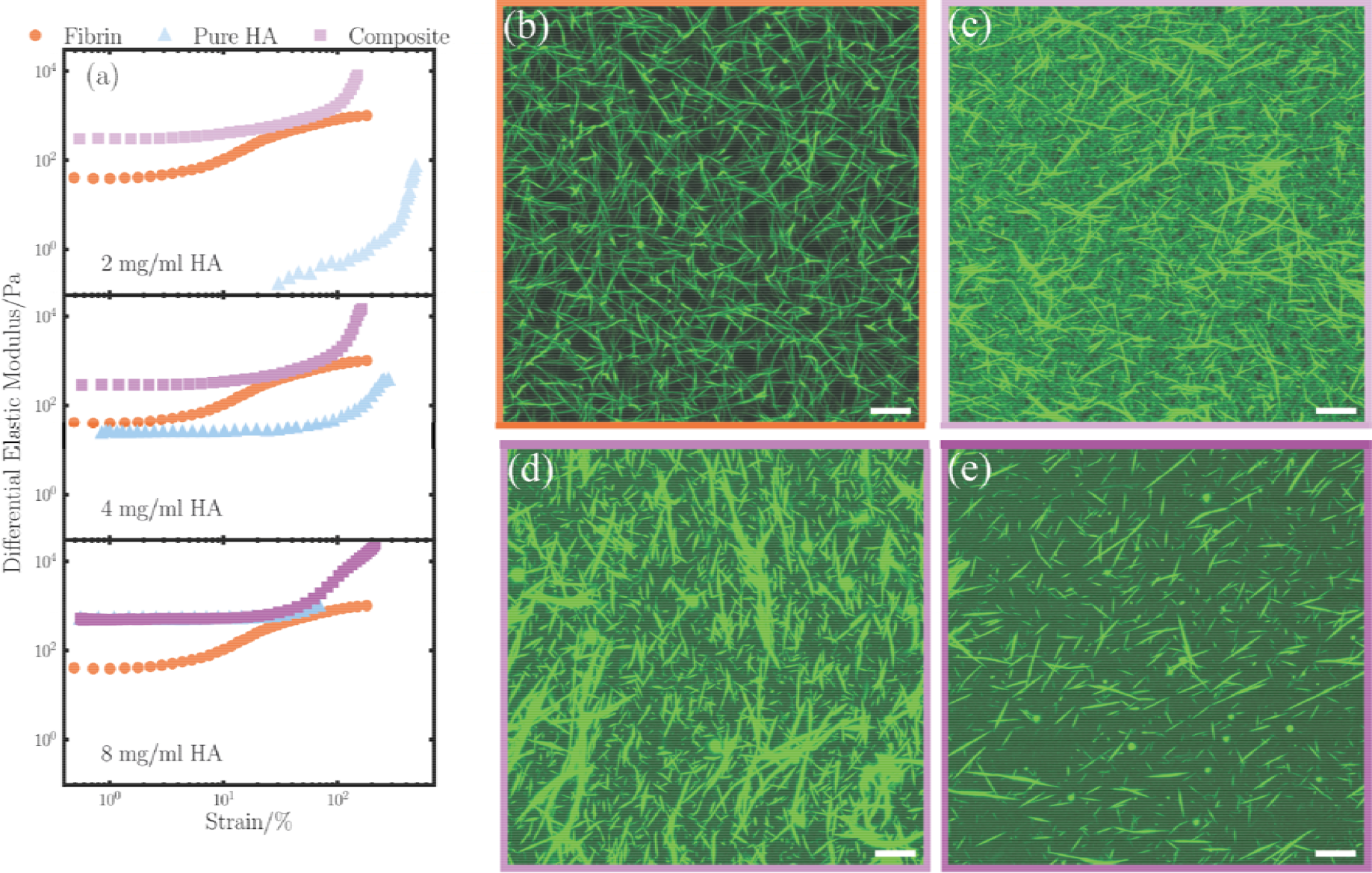
Nonlinear elastic properties and structure of composite hydrogels and pure tyramine modified hyaluronic acid (Tyr-HA) and fibrin. (a) Comparison between the non-linear elastic response of pure Tyr-HA (Δ), pure fibrin (o), and Tyr-HA/fibrin composites (⍰) with a fixed fibrin concentration of 2 mg/ml and varying Tyr-HA concentrations of 2 mg/mL, 4 mg/mL, and 8 mg/mL. Each curve is the mean of 3 repeats. (b-e) Confocal images of 2 mg/mL fibrin networks in absence of Tyr-HA (b), and in composites with 2 mg/ml Tyr-HA (c), 4 mg/ml Tyr-HA (d), and 8 mg/ml Tyr-HA. (e). Images are maximum projections taken over a depth of 14 μm with a spacing of 0.33 μm. Scale bars are each 10 μm.

To assess whether and how the presence of the Tyr-HA network influences the formation of fibrin fibres in the composite materials, we performed confocal fluorescence microscopy. The fibrin network was visualised by addition of fluorescent fibrinogen monomers, the precursor to fibrin, before polymerising the system. We observed clear fibre formation in each of the fibrin-containing samples (Figure 2(b-e)). Qualitative differences were obvious between the formed fibrin networks, with an apparently shorter fibre length as the Tyr-HA concentration increases. To summarise, we observed a marked increase in the linear modulus of composite systems relative to single component systems for lower concentrations of Tyr-HA. For each Tyr-HA concentration, the composite had a higher modulus at high strains than the single component systems.

### 2.3 Hyaluronan Content Governs Cell Morphology in Composite Hydrogels

To assess how the composition of the pure and composite hydrogels influences cell behaviour, we evaluated the early cellular response. Cell morphology was evaluated after 1 or 3 days of encapsulation in the hydrogels using confocal fluorescence images with stained cytoplasm and actin cytoskeleton. A clearly distinct cell morphology was apparent between the high Tyr-HA content hydrogels and the low Tyr-HA hydrogels. In the low Tyr-HA content hydrogels, the hMSCs took on a spread morphology already from day 1, evidenced by many membrane protrusions (data not shown) which became more pronounced at day 3 (figure 3(a-c)). This was obvious in both the pure Tyr-HA hydrogels (figure 3(a-c)) and in the fibrin-containing composite hydrogels (figure 3 (a-c)). By contrast, cells maintained a round morphology in the high Tyr-HA content hydrogels, both in absence and presence of fibrin (figure 3(a-c)). Concurrently, in the low Tyr-HA content hydrogels we observed clear stress fibres forming in the actin cytoskeleton, while in the high Tyr-HA content hydrogels the actin cytoskeleton appeared primarily cortical (figure 3b). Notably, this morphological distinction between cells in low and high Tyr-HA hydrogels cannot be only attributed to changes in the macroscopic linear modulus because we observed distinct morphologies in composites with similar moduli, and similar morphologies when comparing pure Tyr-HA samples with the accompanying, stiffer composites. Instead, only the concentration of Tyr-HA significantly affected the morphology of the encapsulated cells (Figure 3(d), see supplementary table 2 for detailed statistical analysis). These findings suggest that the degree of confinement that a cell locally experiences, governed by Tyr-HA concentration, can influence the behaviour of encapsulated hMSCs.

**Figure 3:**
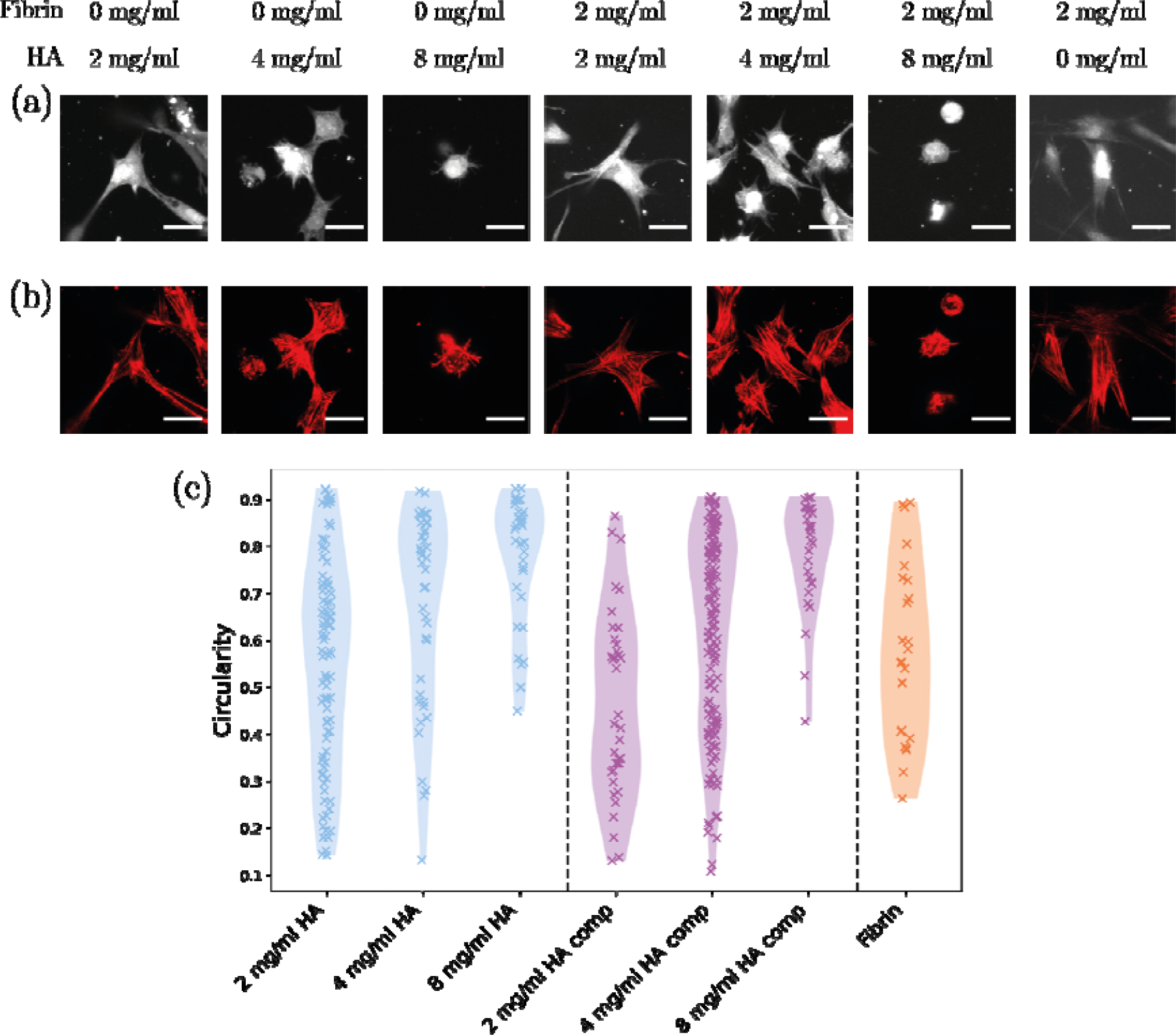
Representative images comparing cell shapes of hMSCs after 3 day culture embedded in pure Tyr-HA hydrogels and composite Tyr-HA/fibrin hydrogels for varying Tyr-HA concentration. (a) Representative images showing the cellular cytoplasm labelled with cFDA. Images are maximum intensity projections of confocal z-stacks over a 20 μm depth and are representative of n=2 experiments and n=3 technical replicates. Scale bars are 30 μm. (b) Corresponding actin signal. (c) Single cell circularity analyses of n>= 25 cells per condition. Data are presented as a violin plot of distribution with each cross representing a single cell. See supp. table 2 for detailed statistics.

### 2.4 Fibrin Governs Cell Shape when Cells are Cultured on Top of Composite Hydrogels

We were surprised that the morphology of cells cultured in composite hydrogels was insensitive to the presence of fibrin, since MSCs express multiple integrin receptors, including αvβ3, α5β1, and αIIbβ3 [33], that can specifically bind to fibrin and facilitate cell adhesion and spreading. We hypothesise that the dense Tyr-HA gel in the pores of the fibrin network may confine the cells and prevent them from adhering to fibrin so they acquire a round morphology. To test the binding capacity of the cells to Tyr-HA and fibrin, we performed 2D culture experiments where we seeded cells on top of the same hydrogel materials instead of embedding them within. Cells seeded on top of Tyr-HA hydrogels maintained a round morphology, implying little to no mechanically active cellular adhesion points (figure 4(a)). This is particularly surprising given that the moduli of the pure Tyr-HA hydrogels span several orders of magnitude. Indeed, despite evidence that CD44 is intracellularly connected to the actin cytoskeleton [34], our results indicated that the adhesions have minimal effect on the cell morphology and actin organisation (figure 3(b)). In contrast, cells seeded on any hydrogel containing fibrin took on a highly spread morphology characterised by long membrane protrusions, implying mechanically active adhesion points (figure 4(a)). Image analysis showed that the circularity of the cells was significantly lower for cells on top of composite hydrogels compared to pure Tyr-HA hydrogels (Figure 4(b), p-value < 0.0001, see supplementary table 3 for detailed statistics). Our findings confirm our hypothesis that fibrin-induced cell spreading is prevented in composite Tyr-HA/fibrin hydrogels by the presence of the dense Tyr-HA meshwork. Our observations suggest that on top of hydrogels, the biochemical nature of the binding substrate determines cell adhesion and spreading, while inside hydrogels, the degree of confinement plays a dominant role.

**Figure 4:**
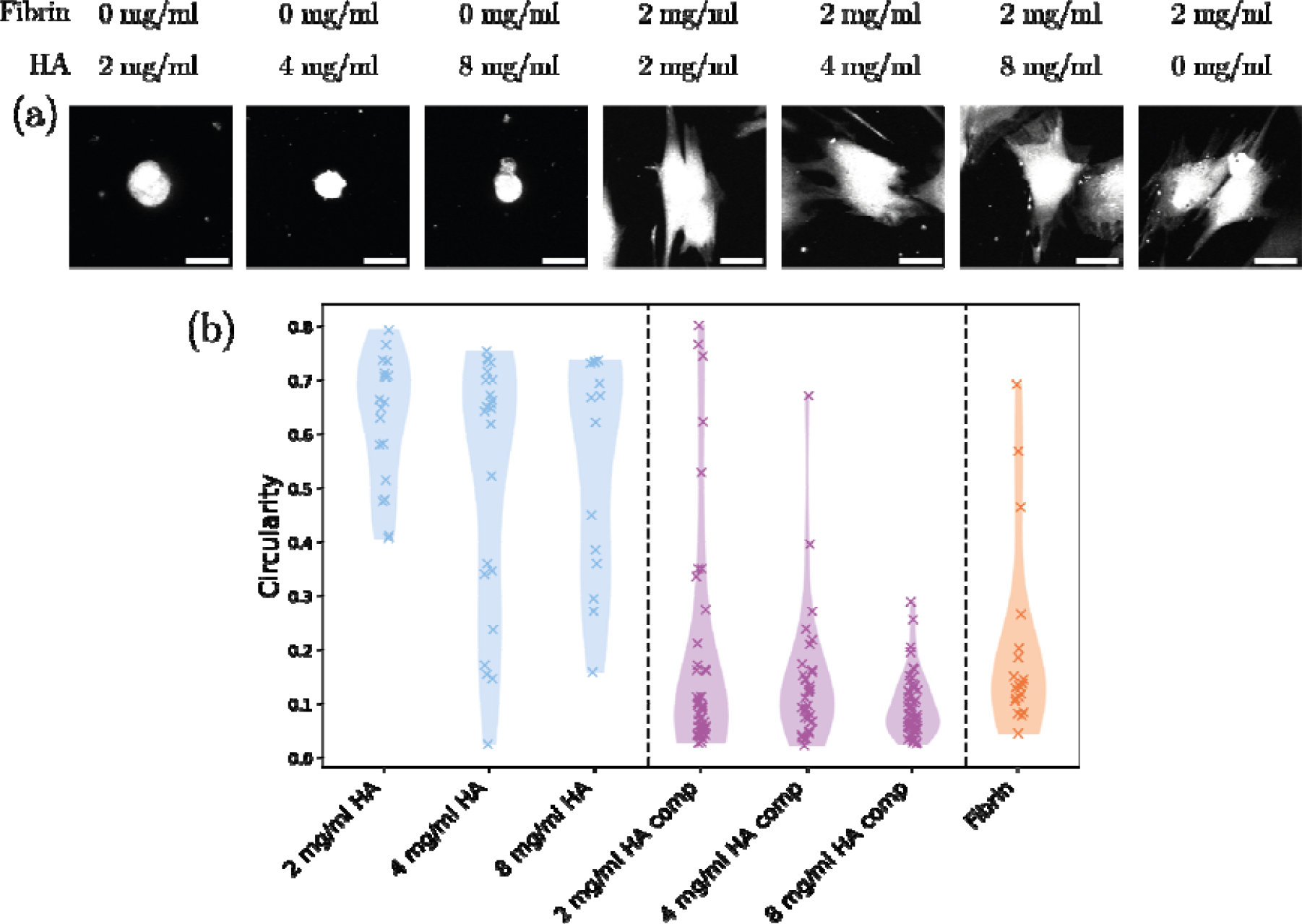
Representative images comparing cell shapes of hMSCs after 3 day culture on the surface of pure Tyr-HA hydrogels and composite Tyr-HA/fibrin hydrogels for varying Tyr-HA concentration. (a) The cytoplasm of the cells is labelled with cFDA. Images are maximum intensity projections of confocal z-stacks over a depth of 20 μm. Scale bars are 30 μm. Images are representative of n = 2 experiments and n = 3 technical replicates. (b) Single cell circularity analyses of n >= 15 cells per condition. Data are presented as violin plots of the distribution with each dot representing a single cell. See supp. table 3 for detailed statistics.

### 2.5 Chondrogenic Genes are Higher Expressed in Composite Hydrogels with More Tyr-HA and Less Fibrin

Finally, we evaluated the potential for hMSCs to differentiate towards the chondrogenic lineage, as cartilage is well suited as a target for tissue engineering using hydrogels and chondrogenic differentiation has been associated with both cell morphology and the stiffness of the environment [35–37]. In each of the hydrogels, hMSCs expressed chondrogenic marker genes *SOX9* (figure 5(a)), *COL2A1* (figure 5(b), *PRG4* (figure 5(c)) and *ACAN* (figure 5(d)) after three days of culture. The presence of sulfated GAG in the hydrogels after 2 weeks of culture also confirmed chondrogenic differentiation (figure 6). We observed significant differences in expression of chondrogenic genes based on the presence of fibrin in the composite materials. Cells in pure Tyr-HA constructs had higher expression of *SOX9, COL2A1* and *PRG4* after 3 days of culture and accumulated a higher content of sulfated GAG per cell during 2 weeks of culture than their respective composite materials. Expression of *ACAN* was higher in both pure and composite constructs containing Tyr-HA in respect to the pure fibrin control. Furthermore, in pure Tyr-HA constructs, expression of *SOX9* and *COL2A1* was higher in cells encapsulated in 2 mg/ml Tyr-HA than 8 mg/ml Tyr-HA constructs (p-value= 0.0180 for *SOX9*, p-value = 0.0139 for *COL2A1*) (figures 5(a-b)). In conclusion, constructs made of pure fibrin gels had the lowest levels of chondrogenic markers and sulfated GAG production, while pure Tyr-HA gels had the highest. The composite materials demonstrated intermediate levels of chondrogenic markers and sulfated GAG content, suggesting that the fibrin/Tyr-HA ratio significantly impacts chondrogenesis.

**Figure 5).**
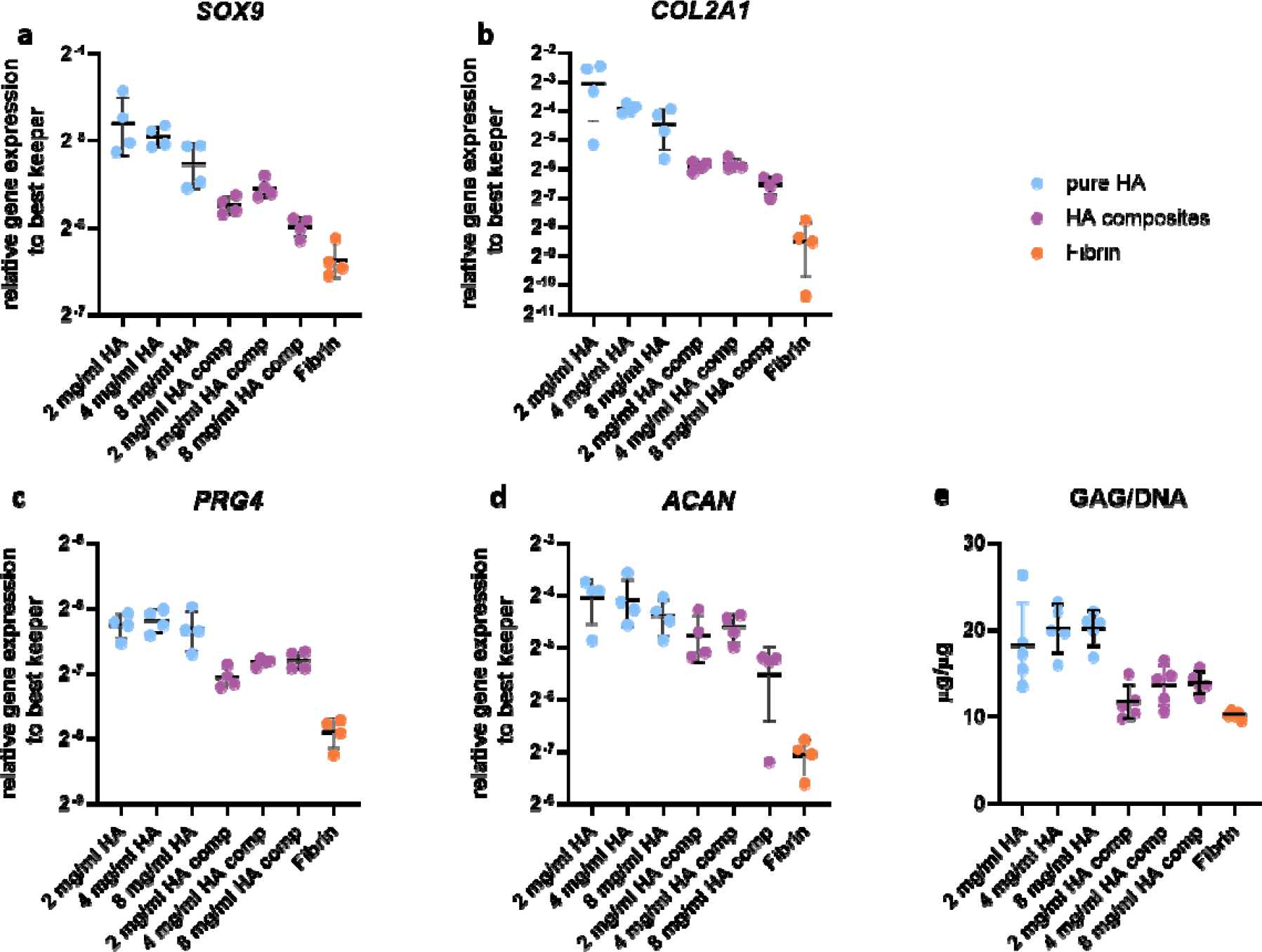
Chondrogenic differentiation of MSCs encapsulated in composite and pure Tyr-HA and fibrin hydrogels. After 3 days of culture, mRNA of chondrogenic genes. (a) *SOX9*, (b) *COL2A1*, (c) *PRG4*, and (d) *ACAN*, each normalised with best keeper genes calculated as geometric mean of the expression of housekeeper genes *GAPDH, UBC* and *RPS27A*. Data are presented as the mean and standard deviation of 4 samples, with each data point representing a separate sample. (e) MSCs encapsulated in pure and composite materials were cultured for 14 days and deposition of GAG was analysed with 1,9-Dimethylmethylene-Blue (DMB) assay and normalised for DNA content evaluated by the CyQUANT Cell Proliferation Assay kit. Data are presented as mean and standard deviation with each dot representing a sample.

## 3. Discussion

In this study, we explored the impact of composite hydrogels made of hyaluronan and fibrin on cell shape and chondrogenic differentiation of mesenchymal stromal cells in the context of cartilage tissue engineering. The materials which we used have been reported separately and in combination [7, 23, 27] to promote chondrogenesis, however the distinctive roles of gel mechanics, structure and biochemistry are unclear. To elucidate these roles we have engineered complex composite materials characterised by enhanced mechanical properties. We found that cell morphologies in the composite materials are primarily governed by the hyaluronan network. Despite evidence of cellular adhesion to fibrin substrates, the presence of fibrin has a minimal effect on morphologies of cells embedded in the composite gels. We attribute this effect to a physical confinement from the local hyaluronan network. Meanwhile, fibrin dominates the macroscopic mechanical response and is able to provide mechanical stability against cell-mediated contraction. Thus, by using composite materials, we can separately control bulk mechanical properties and cellular behaviours.

In our composite materials, we used fibrin and tyramine-modified hyaluronan as the basic building blocks. The hyaluronan forms networks with an extended linear elastic regime, while fibrin forms fibrous networks with nonlinear rheological properties such as strain stiffening (c.f. Figure 2) [38]. We observed an enhancement in the linear modulus of the composites, similarly to previous observations of collagen and hyaluronan [26]. We attribute this to the suppression of fibre bending modes in fibrin, leading to probing of the fibre stretching regime at lower strains than in pure fibrin systems. In particular for our composites, we observed that, while the modulus of the pure Tyr-HA networks increased with concentration, consistent with polymer theories[39], in the composites the linear modulus was far less sensitive to the Tyr-HA concentration, implying that the enhancement of the modulus relative to the pure systems arises due the fibrin network, backing up the idea that we are probing fibrin’s stretch regime at low strain.

Despite previous studies considering how the mechanical environment affects cell behaviour [14, 23, 40], there has been little work studying competing effects of multiple networks, which can inhibit cell protrusions through the smaller pore network, but simultaneously encourage cell protrusions through the fibrous protein network. Our work suggests that this competition plays a key role in determining cell response. While we produced hydrogels with naively similar mechanical properties (the composites of 2 mg/ml Tyr-HA, 2 mg/ml Fibrin compared with 4 mg/ml Tyr-HA, 2 mg/ml Fibrin), the cell morphology when cultured inside these hydrogels depended only on the hyaluronan content and not the presence or absence of fibrin. We attribute this effect to the idea that the cell responds more strongly to its local environment, which is governed by the smaller pore network of the hyaluronan, than to the macroscopic gel stiffness. Indeed, we showed that MSCs in hydrogels with high hyaluronan content exhibited an actin cytoskeleton which is more aligned with the external membrane of the cells, while MSCs in hydrogels with low hyaluronan content exhibited an elongated morphology with the formation of stress fibres, agnostic of the presence or absence of fibrin. This highlights the ability of the hydrogel’s mechanical microenvironment to regulate the actin cytoskeleton and potentially influence chondrogenic differentiation. It is worth noting that the use of chondrogenic media containing TGF beta may have influenced the actin cytoskeleton of the cells by inducing actin reorganisation [41, 42]. However, the actin cytoskeleton can also be affected by dimensionality [43], and substrate [44] so it is difficult to disentangle the effect that TGF beta may have had.

Additionally, the dimensionality of the substrate plays a large role in influencing cell behaviour. In our system the effects of dimensionality arise from differences in confinement in 3D which do not affect the 2D situation. We infer that cell spreading in 3D is controlled by the degree of confinement in a dense hydrogel matrix, rather than by the formation of adhesions by comparing cell morphologies in composite materials with pure hyaluronan materials. This contrasts with previous observations of dimensionality on cell spreading, where in 2D cells spread more on stiffer substrates while in 3D cells spread more in softer environments[16]. It remains unclear why cells spread in low Tyr-HA content pure hyaluronan matrices, given that in 2D we see no evidence for adhesions forming with the matrix. The cells may form cell-cell contacts, which would be reflective of pellet culture which has previously been used to facilitate chondrogenesis of MSCs [45]. Alternatively they may quickly produce their own pericellular environment to which they can bind, the production of which may be more restricted in a denser local environment, i.e., at elevated hyaluronan concentrations [46]. In any case it is clear that these cells form more prominent protrusions when the hyaluronan concentration is lower, independent of the presence or absence of fibrin. It should be noted that the existence of direct cross-links between the cell membrane and tyramine groups has been previously reported, which has an effect on the mechanotransduction in hMSCs [47], but we see minimal effects of this in our cell morphology or gene expression data when considering different Tyr-HA concentrations. The evidence collected in both 2D and 3D environments underscores the importance of considering the 3D microenvironment when developing strategies for cell differentiation and tissue engineering.

In addition to directly observing cellular morphology, we considered cell fate, which can also be controlled by the local environment and may be linked to the cell morphology [28]. Cell differentiation is a complex process that is regulated by a combination of mechanical and biochemical cues, with the mechanical microenvironment playing a critical role in the process. In the context of chondrogenic differentiation of mesenchymal stromal cells (MSCs), the use of hydrogels with varying hyaluronan content can be used to control the mechanical microenvironment [48]. While previous work has found a link between cell shape and eventual cell fate [49], we saw that in composite materials the primary factor governing the expression of chondrogenic genes is from the biochemical makeup of the extracellular matrix. To achieve the highest level of expression in typical chondrogenic genes (*SOX9, ACAN, COL2A1, PRG4*), cells were cultured in pure Tyr-HA matrices, where the evidence in 2D culture implies minimal adhesions. However, in the context of tissue engineering, we argue that a compromise is beneficial, whereby the fibrin network provides structural support to the tissue as a whole, preventing macroscale contraction and enhancing the modulus of the constructs although it leads to lower expression of chondrogenic genes and production of sulfated GAG. Although pellet culture can achieve chondrogenesis [45], we argue, given the low cell density in native articular cartilage, that it is relevant that we can achieve chondrogenesis in biofabricated constructs with a low cell density maintained by resistance to cell-mediated contraction. From a modelling perspective, a lower cell density better reflects the situation in mature articular cartilage, and allows easier study of cell-matrix interactions with minimal influence from cell-cell interactions.

Similarly to our results, previous studies have consistently revealed contrasting effects of fibrin and HA on chondrogenesis. Specifically, fibrin has been shown to exert an inhibitory effect on the differentiation of mesenchymal stem cells (MSCs) towards the chondrogenic lineage [50]. In contrast, HA has emerged as a promising enhancer of chondrogenesis, with studies highlighting its potential to improve the expression of chondrogenic marker genes and promote GAG deposition [51, 52]. Moreover, these studies emphasise the importance of considering the composition and concentration of HA and HA composite materials, as well as the presence of integrin binding sites, as key factors that modulate the efficacy of chondrogenesis in MSCs.

Our work provides valuable insights into the behaviour of cells in response to environmental mechanical stimuli. However, the study has some limitations that should be considered when interpreting the results. Firstly, we only used one type of cell in our experiments, and it is possible that different cell types such as articular chondrocytes, satellite cells or hepatic stem cells may have different responses to the mechanical stimuli. Secondly, we did not explore the effect of other extracellular matrix components such as collagens. Finally, we use specific concentrations of each component, and therefore we can only draw conclusions based on these concentrations, particularly as the composite networks have non-linear behaviours. With these limitations in mind we must be careful when considering our results in terms of the generality to other systems. However, we argue that the fibrin network plays a mostly structural or mechanical role in these constructs, where the cellular behaviour is mostly affected by the hyaluronan network and therefore we expect our results to be applicable to other composites of hyaluronan and fibrous protein networks. Additionally, our results provide valuable mechanistic insights into the combined effects of fibrin and hyaluronan, two clinically utilised biomaterials. By elucidating the intricate interplay between these components within the context of tissue engineering and regenerative medicine, our findings hold promise for unravelling novel mechanistic pathways and informing the design of advanced biomaterials with enhanced therapeutic potential [53].

Our findings highlight the potential of multi-component hydrogels for controlling cellular behaviour and bulk mechanical properties of cell-hydrogel constructs independently, which may aid novel therapeutic strategies for cartilage regeneration. The use of primary human cells and clinically and biologically relevant materials, such as fibrin and hyaluronan, further increases the translational potential and relevance of the findings. By adjusting the relative content of biologically relevant materials, we can design complex hydrogels with improved mechanical properties and stability, while also influencing cell behaviour. Our work also highlights the importance of understanding the complex interplay between mechanical and biochemical cues of the local niche in regulating cell differentiation during development and in disease.

## 4. Acknowledgments

We acknowledge funding from the Convergence programme Syn-Cells for Health(care) under the theme of Health and Technology. This work was partially funded by Open Mind Convergence, the project Medical Delta RegMed4D and the project ‘How cytoskeletal teamwork makes cells strong’ with project number VI.C.182.004 of the NWO Talent Programme which is financed by the Dutch Research Council (NWO). MD acknowledges funding from the European Union’s Horizon 2020 Research and Innovation Programme under Grant Agreement No.874790 (cmRNAbone). We thank Janneke Witte-Bouma and Dr. Eric Farrell from the Department of Oral and Maxillofacial Surgery, Erasmus MC, University Medical Center Rotterdam, the Netherlands, for contributing to the isolation and expansion of cells. We are also grateful to Wendy J L M Koevoet from the department of Otorhinolaryngology, Erasmus MC, University Medical Center Rotterdam, the Netherlands, for her support in biochemical and histological analyses. The authors would like to thank Gert-Jan Kremers from Erasmus MC Optical Imaging Center for his support in confocal imaging. The authors acknowledge Maria Kalogeropoulou from the Department of Bionanoscience, TU Delft for preliminary rheology experiments on composite materials, and Mr Nicolas Devantay for supporting the preparation of the tyramine derivative of hyaluronan.

## Contribution

**M. Fenu**: Conceptualisation, Methodology, Validation, Formal analyses, Writing, Visualisation and Supervision; **I. Muntz**: Conceptualisation, Methodology, Validation, Formal analyses, Writing, Visualisation and Supervision; **D. Harting**: Investigation; **J. Xu**: Investigation; **M. D’este**: Methodology, Investigation, Resources, Supervision; **G.H. Koenderink**: Conceptualisation, Resources, Writing, Supervision, Project Administration, Funding acquisition; **G.J.V.M van Osch**: Conceptualisation, Resources, Writing, Supervision, Project Administration, Funding acquisition.

## Materials and Methods

### Hydrogel Preparation

Tyramine modified lyophilised Hyaluronic Acid (Tyr-HA) (Loebel et al 2015) was hydrated in M-buffer (109 mM sodium chloride; Sigma Aldrich, 20 mM HEPES - N-(2-Hydroxyethyl)piperazine-Nlll-(2-ethanesulfonic acid); Sigma Aldrich, 5.36 mM potassium chloride; Sigma Aldrich - brought to pH 7.3 with 1 M sodium hydroxide) in the presence of horseradish peroxidase (HRP) (stored at 1000 U/ml at -20 °C, thawed on the day of use and kept on ice; Sigma Aldrich) at a final concentration of 2 U/mL. Pure Tyr-HA hydrogels were obtained by cross-linking Tyr-HA solutions with hydrogen peroxide (Sigma Aldrich) at a final concentration of 300 μM. To ensure adequate mixing of the sample, the volumes of the precursor Tyr-HA and hydrogen peroxide solutions were of the same order, with 70 vol% Tyr-HA solution and 30 vol% hydrogen peroxide solution. Samples were mixed by repeatedly pipetting the sample up and down at least 20 times.

For the preparation of the composite systems, human fibrinogen (Enzyme Research Laboratories, plasma purified, plasminogen, von Willebrand factor, fibronectin depleted, FIB 3 5120AL, hydrated, dialysed into M-buffer, flash frozen in liquid nitrogen and stored in 50 μl aliquots at -80 °C) was added to the hydrated Tyr-HA to reach a concentration of 2 mg/mL. Calcium chloride was added to reach a concentration of 5 mM and the sample was left to incubate at 37 °C for 10 minutes. Human thrombin (Enzyme Research Laboratories, HT 5146 L, hydrated to 1000 U/ml, flash frozen in liquid nitrogen and stored in 2.5 μl aliquots at -80 °C) was then added to reach a final concentration of 0.5 U/mL, which initiates the conversion of fibrinogen to fibrin monomers and the subsequent formation of a fibrin network. Within seconds of adding thrombin, hydrogen peroxide was added in the same way as for the pure Tyr-HA networks. The final fibrin concentration was kept constant at 2 mg/mL, while the Tyr-HA concentration was varied between 2 mg/mL, 4 mg/mL or 8 mg/mL.

### Mechanical characterisation

To assess the macroscopic mechanical properties of the hydrogels, shear rheology was performed using a stress-controlled rotational rheometer (Anton Paar MCR 501) equipped with a stainless steel cone plate geometry with a 30 mm diameter and 1° angle. The rheometer was prewarmed to 37 °C and all experiments were performed at 37 °C in a humid environment. Directly after sample preparation, a 130 μl volume of sample was pipetted onto the bottom plate and the rheometer head was brought to the measuring position. The boundary of the sample was coated in mineral oil (lot nr # MKCK4437, Sigma) to prevent solvent evaporation. The samples were allowed to polymerise for 2 hours in order to ensure a steady-state, as monitored by small amplitude oscillatory shear tests during polymerisation with a frequency of 1 Hz and small strain amplitude of 0.5%.

Once the storage modulus reached a plateau, we measured the differential elastic modulus by performing a stress ramp test. The shear stress was increased logarithmically from 0.01 Pa to 10 kPa and the corresponding strain was measured. We measured 10 points per decade of stress with 5 s per point, to ensure each point was measuring a steady state value. The fibrin rheology is independent of the stress rate due to its highly elastic nature [30]. We assume that the hyaluronan rheology is also independent of the stress rate, however, that assumption does not hold for the softest pure hyaluronan gel as we measure a highly viscoelastic response, which can be seen in the frequency dependence of the storage and loss moduli during a frequency sweep (supplementary figure 3). The differential elastic modulus was then calculated from the stress/strain curve through

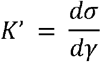

using the gradient function in numpy.

### Structural characterisation

Hydrogel samples were imaged using a Leica Stellaris 8 Falcon confocal microscope with environmental control to maintain a temperature of 37 °C. We used a 63x glycerol immersion objective with a numerical aperture of 1.3. By utilising the digital zoom, the pixel size was set to 100 nm, approximately twice the optimal resolution governed by diffraction. To visualise the fibrin network, fibrinogen conjugated with Alexa Fluor 546 (Thermofisher Scientific) was mixed with non-fluorescent fibrinogen in a mass ratio of 1:19 before polymerizing as described above.

### Primary Cell isolation and expansion

Human mesenchymal stromal cells (hMSC) were isolated from the surplus bone chips from the iliac crest of 4 donors, (10-12-year-old boys and 1 not defined) undergoing alveolar bone graft surgery. Cells were obtained under the approval of the Erasmus MC Ethics Committee (MEC-2014-16). MSC were isolated by plastic adherence and one day after seeding, the non-adherent cells were washed with PBS supplemented with 1% v/v foetal bovine serum (FBS). The cells were expanded at a seeding density of 2300 cells/cm^2^ in Alpha Minimal Essential Medium (α-MEM), supplemented with 10% v/v FBS, 1.5 µg/mL fungizone, 50 µg/mL Gentamicin (all from Thermo Fisher Scientific, Welthon, MA, USA), 25 µg/mL L-ascorbic acid 2-phosphate (Sigma Aldrich, St. Louis, MO, USA), and 1 ng/mL fibroblast growth factor 2 (Instruchemie, Delfzjil, Netherlands) in a humidified atmosphere at 37 °C with 5% CO_2_. Medium was renewed twice or three times a week and cells were trypsinised between 80% and 90% confluency. For the following experiments cells were used between passages 3 and 5.

### Cell seeding and encapsulation

After trypsinisation, hMSCs were washed once with Phosphate Buffered Saline (PBS, Thermo Fisher Scientific, Welthon, MA, USA) and resuspended at different concentrations. To visualise cell morphology, after trypsinisation hMSCs were labelled with CFDA-SE (Vybrant CFDA-SE Cell tracer Kit, Thermo Fisher, Carlsbad, California, United States) following manufacturer instruction at a concentration of 10^6^ cells/ml of PBS/CFDA-SE solution. For 2D experiments, 30 microlitre hydrogels were first crosslinked in 96 well high imaging resolution plates (Greiner Bio-One, Kremsmünster, Austria) and cells were seeded on top at a final density of 5000 cells/ ml (1000 cells/well) in chondrogenic media composed of Dulbecco’s modified Eagle’s medium with Glutamax (DMEM-HG; Gibco, Carlsbad, California, United States) supplemented with 1% Insulin-Transferrin-Selenium (ITS), 1.5µg/mL fungizone,50 µg/mL gentamicin, 1 mM sodium pyruvate (Gibco, Carlsbad, California, United States), 40 µg/mL L-proline (Sigma Aldrich, Missouri, USA), 100nM Dexamethasone (Sigma Aldrich,Missouri, USA), 10 ng/mL recombinant human transforming growth factor beta 1 (TGF-β1 from R&D System, Minnesota, USA). For 3D cultures, cells were resuspended at a density of 15×10^6^ cells/ml in chondrogenic medium and were mixed with pure and composite hydrogel mixture as described above at a volume of 20% of total volume to reach a final concentration 3×10^6^ cells/ml of hydrogels. Hydrogel-cell mixtures were crosslinked in Nuclon^TM^ Delta surface 96-well plates (Thermo Fisher, Carlsbad, California, United States) and cultured in chondrogenic media for 1, 3 and 14 days in a humidified atmosphere at 37 °C with 5% CO_2_ and medium was changed 2-3 times per week. The prepared constructs were tested for cell viability using Viability/Cytotoxicity kit (Invitrogen) following manufacturer instruction. Briefly, cells were stained with calcein AM from live cells and Ethidium homodimer-1 to detect dead cells. Samples were imaged 24 hours post-encapsulation with minimal to no cell death observed across varying Tyr-HA concentrations or with the presence/absence of fibrin (supplementary figure 1).

### Fluorescent labelling and Confocal imaging

For live cell imaging, the cell’s actin cytoskeleton was stained with 5 nmol/l of SiR-actin(Spirochrome, Stein am Rhein, Switzerland) in DMEM for 30 minutes in a humidified atmosphere at 37 °C with 5% CO_2_ at 24 and 72 hours. Afterwards, samples were washed 2 times in PBS and placed afterwards in standard chondrogenic media. Live confocal microscopy was performed with a Leica SP5 confocal microscope (Thermo Fisher Scientific, Welthon, MA, USA) in a humidified atmosphere at 37 °C with 5% CO_2_ either with 20X dry lenses or with 40X oil immersion lenses in bright field, FITC and far red channels. z-stack over a total z-range of 20 μm were acquired with 2 μm z-spacing.

### Image analyses

Confocal stacks were converted to maximum projections in FiJi [54]. Maximum projections were first denoised with a total variation denoising procedure (skimage.restoration.denoise_tv_chambolle) and binarised using an otsu threshold (skimage.filters.threshold_otsu). Cells were then identified using the label function in skimage (skimage.measure.label) which identifies regions in a binary image. Regions were filtered to discard any regions smaller than 140 μm^2^ (equivalent to 1000 pixels or a circle of radius 6.8 μm). It was qualitatively checked that this filter did not remove anything that was identified as a cell by eye. The skimage function regionprops (skimage.measure.regionprops) was used to extract the area and perimeter of each region. The cell morphology in 2D and 3D was quantified by the dimensionless circularity, defined as:

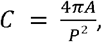

where *A* is the projected area of the cell and *P* is the projected perimeter of the cell. By this metric, a perfect circle has *C* = 1 while a highly spread cell will have *C* → 0. This was calculated for maximum *z*-projections of the captured confocal stacks described above in both 2D and 3D.

### Contractility

Cells encapsulated in hydrogels were cultured for 14 days and afterwards photographs were taken with a mobile phone from the same sample-to-lens distance with a ruler in the image to ensure accurate scale. To evaluate shrinkage, we manually measured in FiJi the diameters of each construct five times, with a total of five constructs per condition.

### Gene expression

Following 3 days of culture, cells encapsulated in hydrogels were harvested and snap-frozen in liquid nitrogen. The constructs were homogenised using a dismembrator (Mikro-Dismembrator S; Sartorius, Goettingen, Germany) and resuspended in RNA STAT-60 REAGENT (RNA-STAT-60; Gentaur, Kampenhout, Belgium). Total RNA was isolated according to the manufacturer’s instructions using the RNeasy Micro Kit (Merk, New Jersey, USA). The concentration and quality of the isolated RNA were measured using the NanoDrop ND100 UV-VIS spectrophotometer (Isogen Life Science B.V, de Meern, the Netherlands). For cDNA preparation, the RevertAid First Strand cDNA Synthesis Kit (ThermoFisher, Carlsbad, California, United States) was used following the manufacturer’s guidelines. The expression of SRY-Box Transcription Factor 9(*SOX9*), collagen type 2 (*COL2A1*), proteoglycan 4 (*PRG4*) and aggrecan (*ACAN*) was assessed through qRT-PCR, which was performed in 10 µL reactions on the ABI Prism 7000 system (Applied Biosystem, Foster City, CA, USA), utilising either Taqman Universal PCR mastermix (Applied Biosystem, Foster City, CA, USA) or SYBR Green (Eurogenetc, Seraing, Belgium). To determine the best housekeeper index (BKI), glyceraldehyde-3-phosphate dehydrogenase (*GAPDH*), Ubiquitin-C (*UBC*) and ribosomal protein S27a (*RPS27a*) were measured and BKI was calculated using the formula (Ct^*GAPDH*\*^ Ct^*RPS*27*a*\*^ Ct^UBC^)^⅓^, where Ct is the cycle threshold for the relevant gene. Relative mRNA levels were calculated using the formula 2^−ΔΔCt^, where ΔΔCt is the difference in cycle threshold between the gene of interest and the BKI. Primers and probes are listed in supplementary table 6.

### Biochemical assays

After 14 days of culture, hydrogel-cell constructs were digested with 4 µg/mL hyaluronidase (Sigma-Aldrich, Missouri, United States) in PBS for 2 hours at 37°C, followed by overnight digestion with a proteinase K solution consisting of 1 mg/mL proteinase K (Sigma-Aldrich, Missouri, United States), 1 mM iodoacetamide (Sigma-Aldrich, Missouri, United States), 10 µg/mL pepstatin A (Sigma-Aldrich, Missouri, United States), 50 mM tromethamine (Tris; Sigma-Aldrich, Missouri, United States) and 1 mM ethylenediaminetetraacetic acid disodium salt dihydrate (EDTA; Sigma-Aldrich, Missouri, United States) buffer pH 7.6. Once completely digested, samples were boiled for 10 minutes to inactivate digestion enzymes. The DNA content of the samples was measured using the CyQUANT Cell Proliferation Assay kit (ThermoFisher, Carlsbad, California, United States) following the manufacturer’s instructions and using the Spectramax iD3 (Molecular Devices, California, United States). To quantify sulfated GAG content, a 1,9-Dimethylmethylene-Blue (DMB) assay was performed. DMB solution is composed of 0.004 g of DMB (AppliChem GmbH, Darmstadt, Germany) to 1.25 mL of 100% EtOH, 0.04 M NaCl, and 0.04 M Glycine and pH was set at 1.75 with pure formic acid. The solution was added to the digested samples. Chondroitin sulfate (Sigma-Aldrich, Missouri, United States) was used as a standard. The solution absorbance was measured at 530 and 590 nm using a VersaMax microplate reader spectrophotometer (Molecular Devices, California, United States) and the concentrations of chondroitin sulfate were calculated as the absorbance at 530 over the absorbance at 590.

### Statistical analyses

Statistical analyses were performed using GraphPad Prism version 8.4.2 (GraphPad Software, San Diego, CA, USA). After graphical representation, the contractility data were analysed to identify outliers using a rout method with a 1% false discovery rate. As a result, one sample of the 4mg/ml Tyr-HA was removed from the following analyses. Normality distributions of the data were tested using the Shapiro-Wilk test. One-Way Ordinary ANOVA with Dunnett’s 3T multiple comparison test was used to determine significant differences between groups. Cell shape analyses were performed using the Kruskal-Wallis test with Dunn’s multiple comparison test. Gene expression data and biochemical data were analysed using the Shapiro-Wilk test to test for normality distribution of the data. One-Way Ordinary ANOVA with Sidak’s multiple comparison test was used to compare differences between groups. The outcomes of statistical analyses can be found in supplementary tables 1, 2, 3, 4, 5.

