## Supplemental Data for "The Impact of the Local Mechanical Environment on Cell Shape and Chondrogenesis of Mesenchymal Stromal Cells in 3D Biomimetic Composite Materials"

**Supplementary Material**

M. Fenu^1,*^, I. Muntz^2,*^, D. Harting^2^, J. Xu^3^, M. D’Este^4^, G.H. Koenderink^2,**^, G.J.V.M van Osch^1,3,5,**^

******* *These Authors contributed equally to the study*

********	*These Authors contributed equally to the study*

***Affiliation***

^1^ Department of Otorhinolaryngology, Erasmus MC, University Medical Center, Rotterdam, The Netherlands

^2^ Department of Bionanoscience, Kavli Institute of Nanoscience Delft, Delft University of Technology, Delft, The Netherlands

^3^ Department of Orthopaedics & Sports Medicine, Erasmus MC, University Medical Center, Rotterdam, The Netherlands;

^4^ AO Research Institute Davos, Davos Platz, Switzerland;

^5^ Department of Biomechanical Engineering, Faculty of Mechanical, Maritime, and Materials Engineering, Delft University of Technology, Delft, The Netherlands

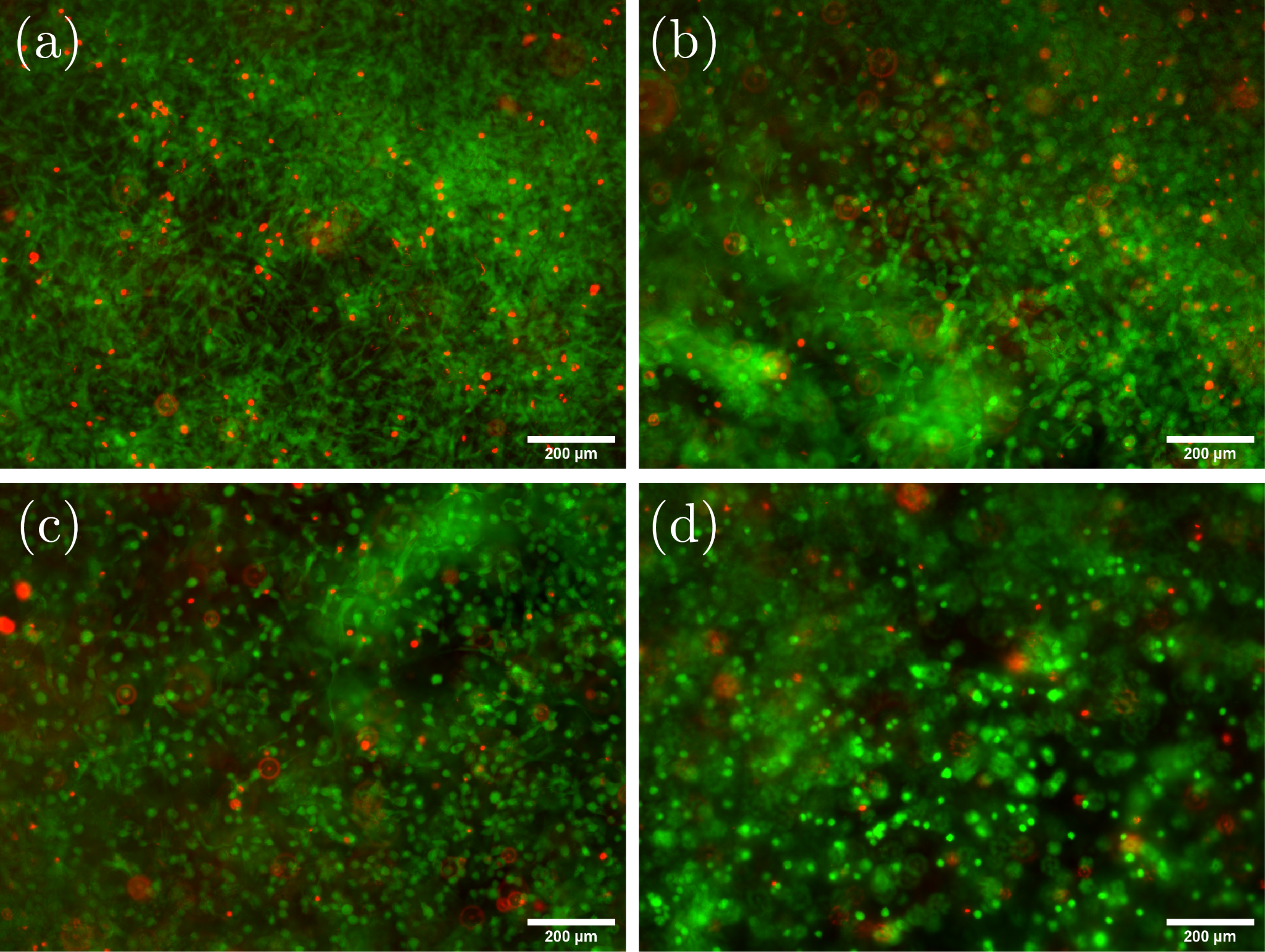

**Supplementary figure 1**: *Representative images of encapsulated MSCs in composites of fibrin and hyaluronic acid after 24 hours*. Live cells are stained in green with calcein AM and dead cell are stained in red with Ethidium homodimer-1. (a) 2 mg/ml Fibrin. (b) 2 mg/ml Fibrin, 2 mg/ml HA. (c) 2 mg/ml Fibrin 4 mg/ml HA. (d) 2 mg/ml Fibrin, 8 mg/ml HA.

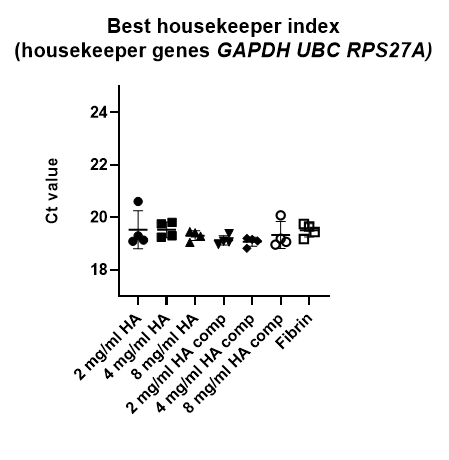

**Supplementary figure 2:** *Best housekeeper index*, calculated as the geometric mean of the mRNA expression of housekeeper genes *GAPDH*, *UBC* and *RPS27A* evaluated with qRT-PCR. Data are presented as the mean and standard deviation of 4 samples, with each *data point* representing a separate sample.

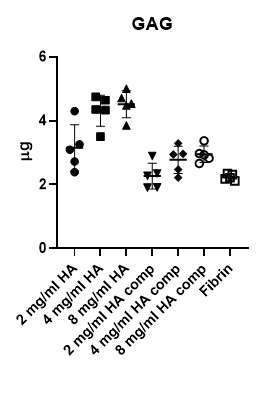

**Supplementary figure 3:** *GAG content of constructs of MSCs encapsulated in pure and composite hydrogels cultured for 14 days in chondrogenic media*. Deposition of GAGs was analysed with DMB assay using Chondroitin Sulphate as standard. Data are presented as mean and standard deviation with each dot representing a sample.

**Supplementary tables**

**Supplementary table 1** *Statistical analyses of contractility assay evaluated with One-Way Ordinary ANOVA.*

| Dunnett's T3 multiple comparisons test | Mean 1 (%) | Mean 2 (%) | Adjusted P Value |
| --- | --- | --- | --- |
| **2 mg/ml HA vs. 4 mg/ml HA** | **21.95** | **63.79** | **<0.0001** |
| **2 mg/ml HA vs. 8 mg/ml HA** | **21.95** | **88.42** | **<0.0001** |
| **2 mg/ml HA vs. 2 mg/ml HA comp** | **21.95** | **99.77** | **<0.0001** |
| **4 mg/ml HA vs. 8 mg/ml HA** | **63.79** | **88.42** | **<0.0001** |
| **4 mg/ml HA vs. 4 mg/ml HA comp** | **63.79** | **100.1** | **<0.0001** |
| **8 mg/ml HA vs. 8 mg/ml HA comp** | **88.42** | **99.58** | **0.0002** |
| 2 mg/ml HA comp vs. 4 mg/ml HA comp | 99.77 | 100.10 | 0.9997 |
| 2 mg/ml HA comp vs. 8 mg/ml HA comp | 99.77 | 99.58 | >0.9999 |
| 2 mg/ml HA comp vs. Fibrin | 99.77 | 100.5 | 0.9386 |
| 4 mg/ml HA comp vs. 8 mg/ml HA comp | 100.1 | 99.58 | 0.9988 |
| 4 mg/ml HA comp vs. Fibrin | 100.1 | 100.5 | 0.9971 |
| 8 mg/ml HA comp vs. Fibrin | 99.58 | 100.5 | 0.9581 |

In bold highlighted significant differences.

**Supplementary table 2** *Statistical analysis of cell morphology in 3D evaluated with* Kruskal-Wallis test*.*

| Dunn's multiple comparisons test | Mean rank 1 | Mean rank 2 | n1 | n2 | Adjusted P Value |
| --- | --- | --- | --- | --- | --- |
| 2 HA vs. 4 HA | 172.4 | 236.0 | 93 | 41 | 0.0528 |
| **2 HA vs. 8 HA** | **172.4** | **295.8** | **93** | **36** | **<0.0001** |
| 2 HA vs. 2 HA/FB | 172.4 | 118.6 | 93 | 37 | 0.2417 |
| 4 HA vs. 8 HA | 236.0 | 295.8 | 41 | 36 | 0.3331 |
| 4 HA vs. 4 HA/FB | 236.0 | 205.9 | 41 | 150 | >0.9999 |
| 8 HA vs. 8 HA/FB | 295.8 | 303.1 | 36 | 30 | >0.9999 |
| **2 HA/FB vs. 4 HA/FB** | **118.6** | **205.9** | **37** | **150** | **0.0008** |
| **2 HA/FB vs. 8 HA/FB** | **118.6** | **303.1** | **37** | **30** | **<0.0001** |
| 2 HA/FB vs. FB | 118.6 | 174.5 | 37 | 25 | 0.8384 |
| **4 HA/FB vs. 8 HA/FB** | **205.9** | **303.1** | **150** | **30** | **0.0005** |
| 4 HA/FB vs. FB | 205.9 | 174.5 | 150 | 25 | >0.9999 |
| **8 HA/FB vs. FB** | **303.1** | **174.5** | **30** | **25** | **0.0008** |

In bold highlighted significant differences.

**Supplementary table 3** *Statistical analysis of cell morphology in 2D evaluated with* Kruskal-Wallis test.

| Dunn's multiple comparisons test | Mean rank 1 | Mean rank 2 | n1 | n2 | Adjusted P Value |
| --- | --- | --- | --- | --- | --- |
| 2 HA vs. 4 HA | 180.8 | 163.0 | 19 | 23 | >0.9999 |
| 2 HA vs. 8 HA | 180.8 | 173.3 | 19 | 15 | >0.9999 |
| **2 HA vs. 2 HA/FB** | **180.8** | **80.28** | **19** | **46** | **<0.0001** |
| 4 HA vs. 8 HA | 163.0 | 173.3 | 23 | 15 | >0.9999 |
| **4 HA vs. 4 HA/FB** | **163.0** | **85.15** | **23** | **34** | **<0.0001** |
| **8 HA vs. 8 HA/FB** | **173.3** | **68.34** | **15** | **53** | **<0.0001** |
| 2 HA/FB vs. 4 HA/FB | 80.28 | 85.15 | 46 | 34 | >0.9999 |
| 2 HA/FB vs. 8 HA/FB | 80.28 | 68.34 | 46 | 53 | >0.9999 |
| 2 HA/FB vs. FB | 80.28 | 108.1 | 46 | 20 | >0.9999 |
| 4 HA/FB vs. 8 HA/FB | 85.15 | 68.34 | 34 | 53 | >0.9999 |
| 4 HA/FB vs. FB | 85.15 | 108.1 | 34 | 20 | >0.9999 |
| 8 HA/FB vs. FB | 68.34 | 108.1 | 53 | 20 | 0.1519 |

In bold highlighted significant differences.

**Supplementary table 4.a** *Statistical analysis* *of gene expression SOX9 evaluated with One-Way Ordinary ANOVA*.

| Sidak's multiple comparisons test | Mean 1 | Mean 2 | Adjusted P Value |
| --- | --- | --- | --- |
| 2 HA vs. 4 HA | 0.03584 | 0.03231 | 0.9357 |
| **2 HA vs. 8 HA** | **0.03584** | **0.02602** | **0.018** |
| **2 HA vs. 2 HA/FB** | **0.03584** | **0.01873** | **<0.0001** |
| 4 HA vs. 8 HA | 0.03231 | 0.02602 | 0.3026 |
| **4 HA vs. 4 HA/FB** | **0.03231** | **0.02150** | **0.0075** |
| **8 HA vs. 8 HA/FB** | **0.02602** | **0.01577** | **0.0123** |
| 2 HA/FB vs. 4 HA/FB | 0.01873 | 0.02150 | 0.9895 |
| 2 HA/FB vs. 8 HA/FB | 0.01873 | 0.01577 | 0.9819 |
| 2 HA/FB vs. FB | 0.01873 | 0.01208 | 0.2369 |
| 4 HA/FB vs. 8 HA/FB | 0.02150 | 0.01577 | 0.4278 |
| **4 HA/FB vs. FB** | **0.02150** | **0.01208** | **0.0256** |
| 8 HA/FB vs. FB | 0.01577 | 0.01208 | 0.9148 |

In bold highlighted significant differences.

**Supplementary table 4.b** *Statistical analysis* *of gene expression COL2A1 evaluated with One-Way Ordinary ANOVA*.

| Sidak's multiple comparisons test | Mean 2 | Mean Diff. | Adjusted P Value |
| --- | --- | --- | --- |
| 2 HA vs. 4 HA | 0.06634 | 0.05404 | 0.1477 |
| **2 HA vs. 8 HA** | **0.04537** | **0.07501** | **0.0139** |
| **2 HA vs. 2 HA/FB** | **0.01688** | **0.1035** | **0.0005** |
| 4 HA vs. 8 HA | 0.04537 | 0.02097 | 0.9874 |
| 4 HA vs. 4 HA/FB | 0.01759 | 0.04875 | 0.2489 |
| 8 HA vs. 8 HA/FB | 0.01085 | 0.03452 | 0.7121 |
| 2 HA/FB vs. 4 HA/FB | 0.01759 | -0.0007 | >0.9999 |
| 2 HA/FB vs. 8 HA/FB | 0.01085 | 0.006032 | >0.9999 |
| 2 HA/FB vs. FB | 0.002782 | 0.01410 | 0.9997 |
| 4 HA/FB vs. 8 HA/FB | 0.01085 | 0.006736 | >0.9999 |
| 4 HA/FB vs. FB | 0.002782 | 0.01481 | 0.9995 |
| 8 HA/FB vs. FB | 0.002782 | 0.00807 | >0.9999 |

In bold highlighted significant differences.

**Supplementary table 4.c** *Statistical analysis* *of gene expression PRG4 evaluated with One-Way Ordinary ANOVA*.

| Sidak's multiple comparisons test | Mean 1 | Mean 2 | Adjusted P Value |
| --- | --- | --- | --- |
| 2 HA vs. 4 HA | 0.01305 | 0.01377 | 0.9997 |
| 2 HA vs. 8 HA | 0.01305 | 0.01254 | >0.9999 |
| **2 HA vs. 2 HA/FB** | **0.01305** | **0.007478** | **0.0002** |
| 4 HA vs. 8 HA | 0.01377 | 0.01254 | 0.9615 |
| **4 HA vs. 4 HA/FB** | **0.01377** | **0.008844** | **0.0009** |
| **8 HA vs. 8 HA/FB** | **0.01254** | **0.009015** | **0.0249** |
| 2 HA/FB vs. 4 HA/FB | 0.007478 | 0.008844 | 0.9193 |
| 2 HA/FB vs. 8 HA/FB | 0.007478 | 0.009015 | 0.84 |
| **2 HA/FB vs. FB** | **0.007478** | **0.00419** | **0.0433** |
| 4 HA/FB vs. 8 HA/FB | 0.008844 | 0.009015 | >0.9999 |
| **4 HA/FB vs. FB** | **0.008844** | **0.00419** | **0.0018** |
| **8 HA/FB vs. FB** | **0.009015** | **0.00419** | **0.0012** |

In bold highlighted significant differences.

**Supplementary table 4.d** *Statistical analysis* *of gene expression* *ACAN evaluated with One-Way Ordinary ANOVA*.

| Sidak's multiple comparisons test | Mean 1 | Mean 2 | Adjusted P Value |
| --- | --- | --- | --- |
| 2 HA vs. 4 HA | 0.06035 | 0.05896 | >0.9999 |
| 2 HA vs. 8 HA | 0.06035 | 0.04768 | 0.8719 |
| 2 HA vs. 2 HA/FB | 0.06035 | 0.03670 | 0.1387 |
| 4 HA vs. 8 HA | 0.05896 | 0.04768 | 0.9374 |
| 4 HA vs. 4 HA/FB | 0.05896 | 0.04105 | 0.4651 |
| 8 HA vs. 8 HA/FB | 0.04768 | 0.02179 | 0.0795 |
| 2 HA/FB vs. 4 HA/FB | 0.03670 | 0.04105 | >0.9999 |
| 2 HA/FB vs. 8 HA/FB | 0.03670 | 0.02179 | 0.7139 |
| **2 HA/FB vs. FB** | **0.03670** | **0.007452** | **0.033** |
| 4 HA/FB vs. 8 HA/FB | 0.04105 | 0.02179 | 0.3629 |
| **4 HA/FB vs. FB** | **0.04105** | **0.007452** | **0.0101** |
| 8 HA/FB vs. FB | 0.02179 | 0.007452 | 0.7596 |

In bold highlighted significant differences.

**Supplementary table 5** *Statistical analysis* *of* *GAG/DNA evaluated with One-Way Ordinary ANOVA*.

| Sidak's multiple comparisons test | Mean 1 | Mean 2 | Adjusted P Value |
| --- | --- | --- | --- |
| 2 HA vs. 4 HA | 18.21 | 20.16 | 0.966 |
| 2 HA vs. 8 HA | 18.21 | 20.16 | 0.9655 |
| **2 HA vs. 2 HA/FB** | **18.21** | **11.68** | **0.0053** |
| 4 HA vs. 8 HA | 20.16 | 20.16 | >0.9999 |
| **4 HA vs. 4 HA/FB** | **20.16** | **13.65** | **0.0055** |
| **8 HA vs. 8 HA/FB** | **20.16** | **13.88** | **0.0079** |
| 2 HA/FB vs. 4 HA/FB | 11.68 | 13.65 | 0.9629 |
| 2 HA/FB vs. 8 HA/FB | 11.68 | 13.88 | 0.9216 |
| 2 HA/FB vs. FB | 11.68 | 10.22 | 0.9967 |
| 4 HA/FB vs. 8 HA/FB | 13.65 | 13.88 | >0.9999 |
| 4 HA/FB vs. FB | 13.65 | 10.22 | 0.4273 |
| 8 HA/FB vs. FB | 13.88 | 10.22 | 0.3371 |

In bold highlighted significant differences.

**Supplementary table 6.** *List of primers used to detect mRNA levels by qRT-PCR. For SYBR green dye based reaction no extra probe is needed as the dye will attach to double strand DNA and therefore detect PCR reaction. For the TaqMan reaction, the TaqMan probe-based qRT-PCR analysis exploits the 5′ → 3′ nuclease activity of the Taq polymerase to detect and quantify specific PCR products as the reaction proceeds.*

| Gene | *Forward primer* | *Reverse Primer* | *Probe* |
| --- | --- | --- | --- |
| *GAPDH* | *5’- ATGGGGAAGGTGAAGGTCG-3* | *5’-TAAAAGCAGCCCTGGTGACC-3’* | *5’-CGCCCAATACGACCAAATCCGTTGAC-3’* |
| *UBC* | *5’-ATTTGGGTCGCGGTTCTTG-3’* | *5’-TGCCTTGACATTCTCGGATGGT-3’* |  |
| *RPS27a* | *5’-TGGCTGTCCTGAAATATTATAAGGT-3’* | *5’- CCCCAGCACCACATTCATCA-3’* |  |
| *PRG4* | *5’- TTGCGCAATGGGACATTAGTT-3’* | *5’-AGCTGGAGATGGTGGACTGAA-3’* |  |
| *ACAN* | *5’-TCGAGGACAGCGAGGCC-3’* | *5’- TCGAGGGTGTAGCGTGTAGAGA-3’* | *5’-ATGGAACACGATGCCTTTCACCACGA-3’* |
| *COL2A1* | *5’-GGCAATAGCAGGTTCACGTACA-3’* | *5’-CGATAACAGTCTTGCCCCACTT-3’* | *5’-CCGGTATGTTTCGTGCAGCCATCCT-3’* |
| *SOX9* | *5’-CAACGCCGAGCTCAGCA-3’* | *5’-TCCACGAAGGGCCGC-3’* | *5’-TGGGCAAGCTCTGGAGACTTCTGAACG-3’* |
